# Cascade screening following a polygenic risk score test: what is the risk of a relative conditional on a high score of a proband?

**DOI:** 10.1101/2021.04.11.439329

**Authors:** Shai Carmi

## Abstract

Polygenic risk scores (PRSs) for predicting disease risk have become increasingly accurate, leading to rising popularity of PRS tests. Consider an individual whose PRS test has placed him/her at the top *q*-quantile of genetic risk. Recently, Reid et al. (Circ Genom Precis Med. 2021;14:e003262) have investigated whether such a finding should motivate cascade screening in the proband’s siblings. Specifically, using data from the UK biobank, Reid et al. computed the empirical probability of a sibling of the proband to also have a PRS at the top *q*-quantile. In this short note, I use the liability threshold model to compute this probability analytically (for either a sibling of the proband or for a more distant relative), showing excellent agreement with the empirical results of Reid et al., including that this probability is disease-independent. Further, I compute the probability of the relative of the proband to be affected, as a function of the quantile threshold *q*, the proportion of variance explained by the score, and the disease prevalence.

## Introduction

Polygenic risk scores show great promise for personalized disease risk prediction. A polygenic risk score (PRS) for a disease is a count of the number of risk alleles carried by an individual, with each allele weighted by its effect size. For complex diseases, a PRS represents the cumulative risk generated by thousands or more variants, each of a small effect^1^. PRSs were empirically shown to explain a substantial proportion of the variance in disease liability, and individuals at the highest PRS percentiles were shown to have risk that can be even ≈3x higher compared to the population mean^2–4^. This suggests the feasibility of personalized prevention and/or intervention in these individuals, whose high risk may not always manifest as traditional risk factors^5^.

In clinical genetics, cascade screening refers to the testing of relatives of individuals who were either found to have a disease with a genetic component or were found to carry a disease mutation^6^. Cascade screening is important, as it allows high-specificity identification of individuals at high genetic risk. Cascade screening is currently limited to severe diseases and single mutations of large effects^7^. However, the development of increasingly accurate polygenic risk scores^8^, along with the increasing number of individuals receiving PRS results worldwide^9^, raise the prospects of performing cascade screening already upon the finding of an extreme PRS.

Recently, Reid et al. have investigated the question of cascade screening following a PRS finding^10^. They empirically studied the following setting. Suppose an individual has taken a PRS test that has placed him/her at the top *q*-quantile of the PRS distribution (where *q* can be, e.g., 1%, 5%, 10%, etc.). What is the probability of a sibling of the proband to also have PRS at the top *q*quantile? Using data on ≈23,000 sib-pairs from the UK biobank, Reid et al. have shown that the siblings of those probands have an increased probability to have a high PRS compared to the population average. Further, the probability was about the same across four diseases.

Despite these interesting results, several questions remain open. Specifically, it is unclear (1) whether the empirical results of Reid et al. (and particularly, the independence across diseases) are concordant with predictions of quantitative genetics theory; (2) what is the expected risk for diseases or threshold quantiles not empirically studied; (3) what is the expected risk for more distant relatives; and (4) what is the expected risk of the relative to become affected, rather than just have a PRS above a cutoff. The latter is important, as the cost-effectiveness of cascade screening depends on the actual disease risk rather than just the PRS. Here, I use the liability threshold model, an established model of disease risk in quantitative genetics, to address these questions.

## Methods

### Model

The liability threshold model (LTM) is a classic model in quantitative genetic theory that relates the risk of a disease to underlying genetic and non-genetic factors^11,12^. According to the LTM, a disease has an underlying continuous liability, distributed as a standard normal random variable. The liability can be written as *y* = *g* + *∈*, where *g* ~ *N*(0, *h*^2^), represents polygenic genetic factors (with variance equal to the heritability *h*^2^) and *∈* ~ *N*(0,1 − *h*^2^) represents non-genetic (environmental) factors. An individual is affected whenever his/her liability exceeds a threshold. For a disease with prevalence *K*, the threshold is *Z*_*K*_ the upper-*K* quantile of the standard normal distribution. (For example, for *K* = 0.01, *Z*_*K*_ = 2.33.) The LTM was found to fit well genetic data on complex diseases, and it is widely applied^13–19^.

In our setting, we do not know the precise value of the genetic factors influencing the liability. Instead, we have an estimate represented by the PRS. We thus write the liability as *y* = *s* + *e*, where *s* is the PRS and *e* is the residual liability, representing non-modeled genetic factors and non-genetic factors^17,19,20^. We assume that the PRS is normally distributed and has been standardized, such that *s* ~ *N*(0, *r*^2^), where *r*^2^ is the variance in liability explained by the PRS. Consequently, *e* ~ *N*(0,1 − *r*^2^).

Consider next a pair of (full) siblings. Using standard quantitative genetic theory, it can be shown that that the PRSs of the two sibs can be written as *s*_1_ = *c* + *x*_1_ for sib 1 and *s*_2_ = *c* + *x*_2_ for sib 2. In these equations, *c* is a genetic component shared between the sibs, equal to the average maternal and paternal PRSs. Its distribution across the population is *c* ~ *N*(0,*r*^2^ /2). Then, *x*_1_ ~ *x*_2_ ~ *N*(0,*r*^2^ /2) are two *independent* genetic components. For details on the derivation, see our previous publications^21,22^. For an intuitive explanation, the “segregation” variance, i.e., the variance of any polygenic component across siblings due to the randomness of meiosis, is known to be half the variance in the population^23^. Thus, given the parental PRSs, the PRS of each child has variance *r*^2^ /2.

We next define a threshold above which an individual is designated as having “high PRS”. We define the threshold as the upper-*q* quantile of the PRS distribution. For example, if *q* = 0.01, an individual is considered to have high PRS if his/her PRS is at the top 1% of PRSs across the population. The PRS has zero mean and variance *r*^2^ in the population, and thus, the value of the PRS at the threshold is *z*_*q*_ *r*, where *z*_*q*_ is the upper-*q* quantile of a standard normal variable. We are told that sib 1 (the proband) has a high PRS, i.e., *s*_1_ > *z*_*q*_ *r*. We would like to compute the conditional probability that sib 2 either also has high PRS, or is affected.

Our calculations are similar to those of So et al.^17^ and Do et al.^19^, who have considered the problem of predicting disease risk based on the PRS of an individual and/or a relative, along with the disease status of the relative. Here, we assume the disease status of the proband is unknown (e.g., for a late-onset disease).

### The probability that the sibling has high PRS

We would like to compute the probability *P*(*s*_2_ > *z*_*q*_ *r* | *s*_1_ > *z*_*q*_ *r*). Using the definition of the conditional probability,

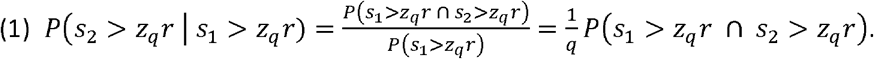

To compute the numerator, we condition on *c*. Given *c*, the scores of the two sibs are independent, i.e.,

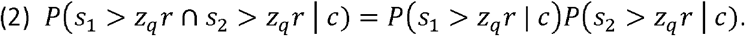

For *i* = 1,2,

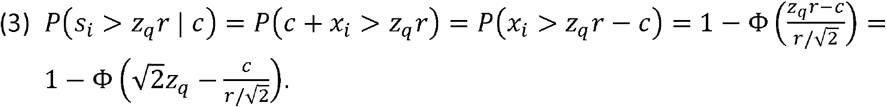

In Eq. (3), Φ(*x*) is the cumulative distribution of the standard normal variable, and we used the fact that *x*_*i*_ ~ *N*(0, *r*^2^ /2). We can now compute the desired probability (Eq. (1)) by substituting Eq. (3) into Eq. (2), and integrating over all *c* ~ *N*(0, *r*^2^ /2),

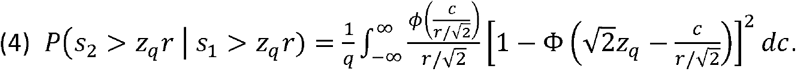

In Eq. (4), *ϕ*(*x*) is the density of the standard normal variable. We now change variables, 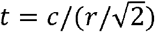, and obtain

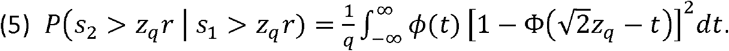

Eq. (5) is our final result for the probability of the sibling of the proband to have high PRS. Note that the probability does not depend on *r*^2^, the variance explained by the PRS, and thus is disease- and PRS-independent.

### The probability that the sibling is affected

Denote by *y*_1_ and *y*_2_ the liabilities of the two sibs, and recall that sib 2 will be affected if *y*_2_ > *z*_*K*_, where *z*_*K*_ is the upper-*K* quantile of the standard normal distribution and *K* is the prevalence. We would like to compute the following probability,

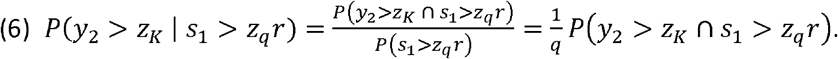

As above, we write *s*_*i*_ = *c* + *x*_*i*_ for *i* = 1,2, and condition on *c*. First, we note that

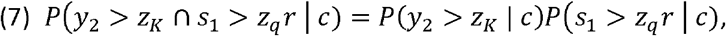

because knowledge of the score of sib 1, given *c*, is not informative on the remaining liability components of sib 2. As in Eq. (3) above,

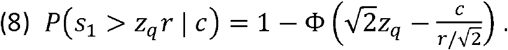

Next,

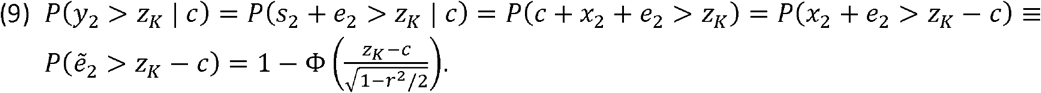

Above, we defined 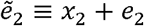 and used the fact that *x*_2_ and *e*_2_ are independent normals, such that 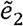 is normal with zero mean and variance 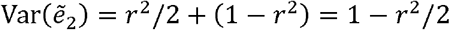. We can compute the desired probability (Eq. (6)) by substituting Eqs. (8) and (9) into Eq. (7), and integrating over all *c*,

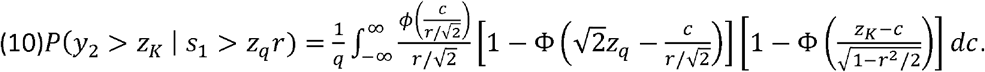

We can again change variables, 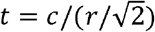, and obtain

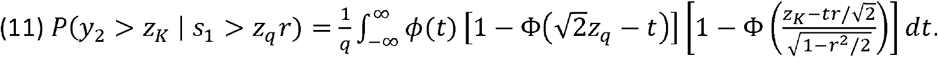

This is our final expression for the probability of the sibling of the proband to be affected. Here, the probability depends on *r*^2^, as well as on the prevalence *K*.

### The probability that an arbitrary relative has high PRS

We now turn from siblings into *d*-degree relatives. [Parents and children and full siblings are first degree relatives; half-sibs, grandparent and grandchild, and uncle and nephew are second-degree relatives; (full) first-cousins are third-degree relatives, and so on.] To compute the probability of the relative to be affected in this more general case, we take a different approach. Denote by *s*_1_ the PRS of the proband and by *s*_2_ the PRS of the relative. It is well known in quantitative genetics^12,21^ that the distribution of (*s*_1_, *s*_2_) is multivariate normal, with the following parameters,

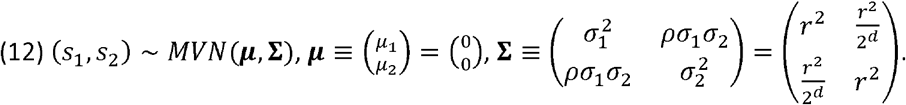

The covariance term is Cov(*s*_1_, *s*_2_) = 2^−*d*^ · *r*^2^ (i.e., the correlation is *ρ* = 2^−*d*^) because 2^−*d*^ is the relatedness coefficient between *d*-degree relatives. As above, we would like to compute the probability *P*(*s*_2_ > *z*_*q*_ *r* | *s*_1_ > *z*_*q*_ *r*) that the relative has high PRS given that the proband has high PRS (top *q*-quantile).

Based on properties of multivariate normal distributions, we have

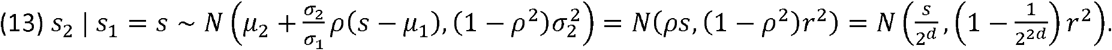

The tail probability of *s*_2_ conditional on *s*_1_ is

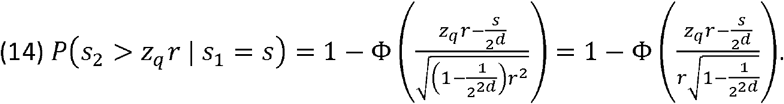

We can now compute the desired probability by integrating over *s*_1_, recalling that *s*_1_ ~ *N*(0, *r*^2^). Denote the probability density function of *s*_1_ as 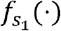. We have

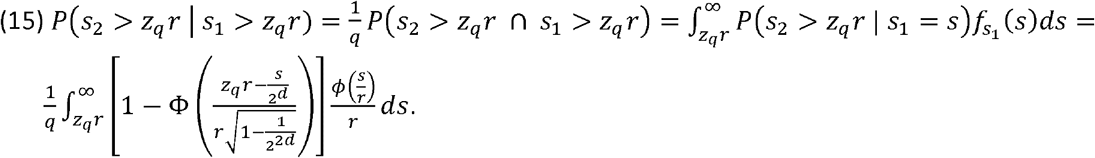

We now change variables, *t* = *s*/*r*, and obtain

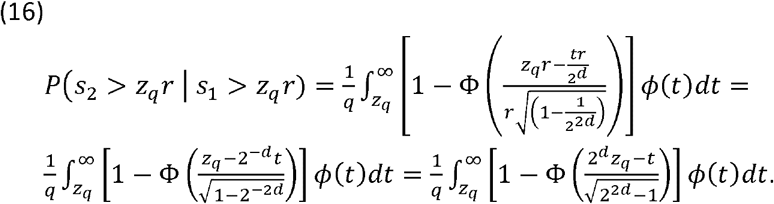

Eq. (16) is our final result for the probability that the *d*-degree relative has high PRS given that the proband has high PRS. We validated numerically that for full siblings (*d* = 1) Eq. (16) gives the same result as Eq. (5). Note that here too, the probability does not depend on *r* and hence is disease- and PRS-independent.

### The probability that an arbitrary relative is affected

Finally, we compute the probability that the relative of the proband is affected. As above, we denote by *y*_1_ and *y*_2_ the liabilities of the proband and relative, respectively, and recall that the relative will be affected if *y*_*z*_ > *z*_*K*_. As in Eq. (15) above,

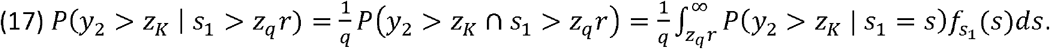

Next,

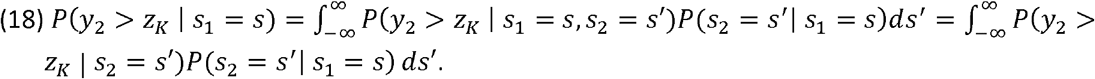

The last step follows because given that we know the PRS of the relative, the total liability (and henceforth the disease status) of the relative no longer depend on the PRS on the proband. Recall that the liability is *y* = *s* = *e*, where *e* ~ *N*(0,1 − *r*^2^). Thus,

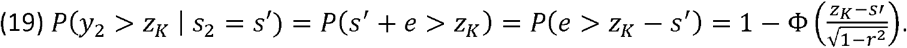

Using Eq. (13) above,

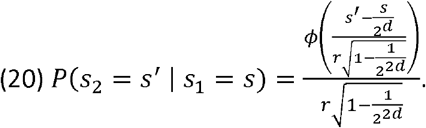

Substituting Eqs. (19) and (20) into Eq. (18) we obtain

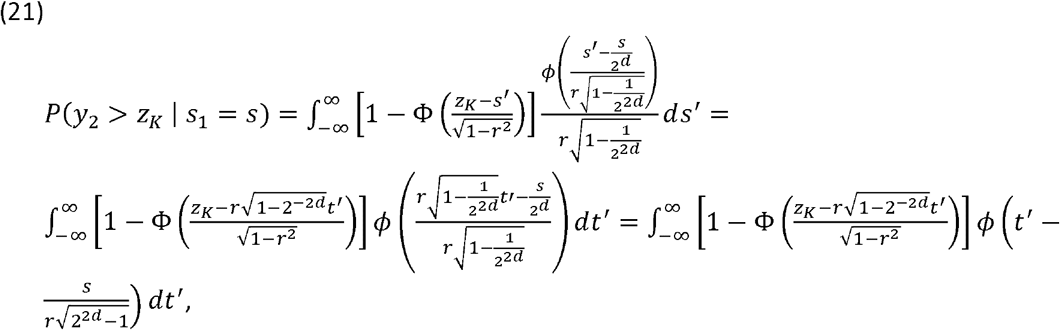

where we changed variables, 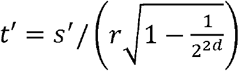. We finally plug Eq. (21) into Eq. (17), recalling again that *s*_1_ ~ *N*(0,1 − *r*^2^). This gives

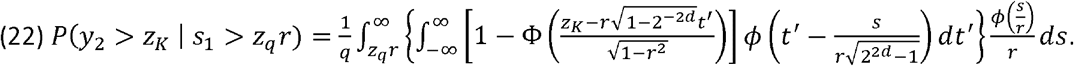

Changing variables, *t* = *s*/*r*, we obtain

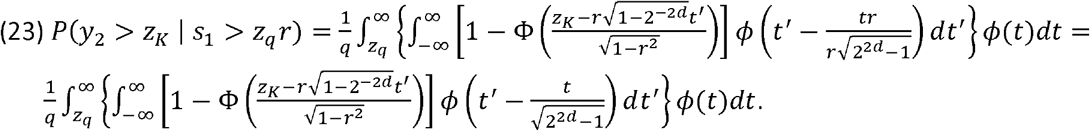

Eq. (23) is our final equation for the probability of the relative to be affected. For the case of full siblings (*d* = 1), we validated numerically that Eq. (23) gives the same result as Eq. (11). As for siblings, the probability of disease depends *r*^2^ and *K*.

### R implementation

For siblings, we solved the integrals in Eqs. (5) and (11) numerically using the function integrate in R. Our code is as follows.

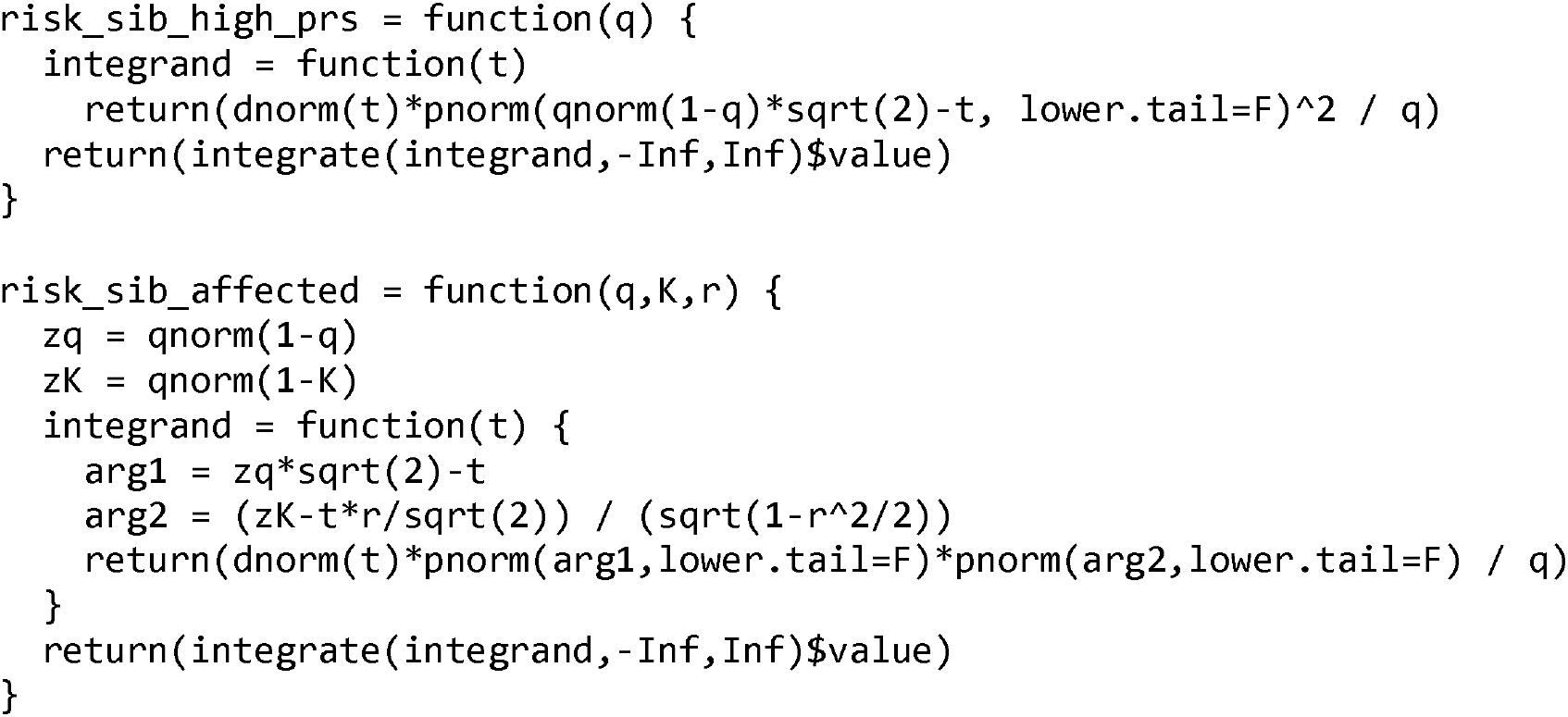

For *d*-degree relatives, we again solved the integrals in Eqs. (16) and (23) numerically in R. Our code is as follows.

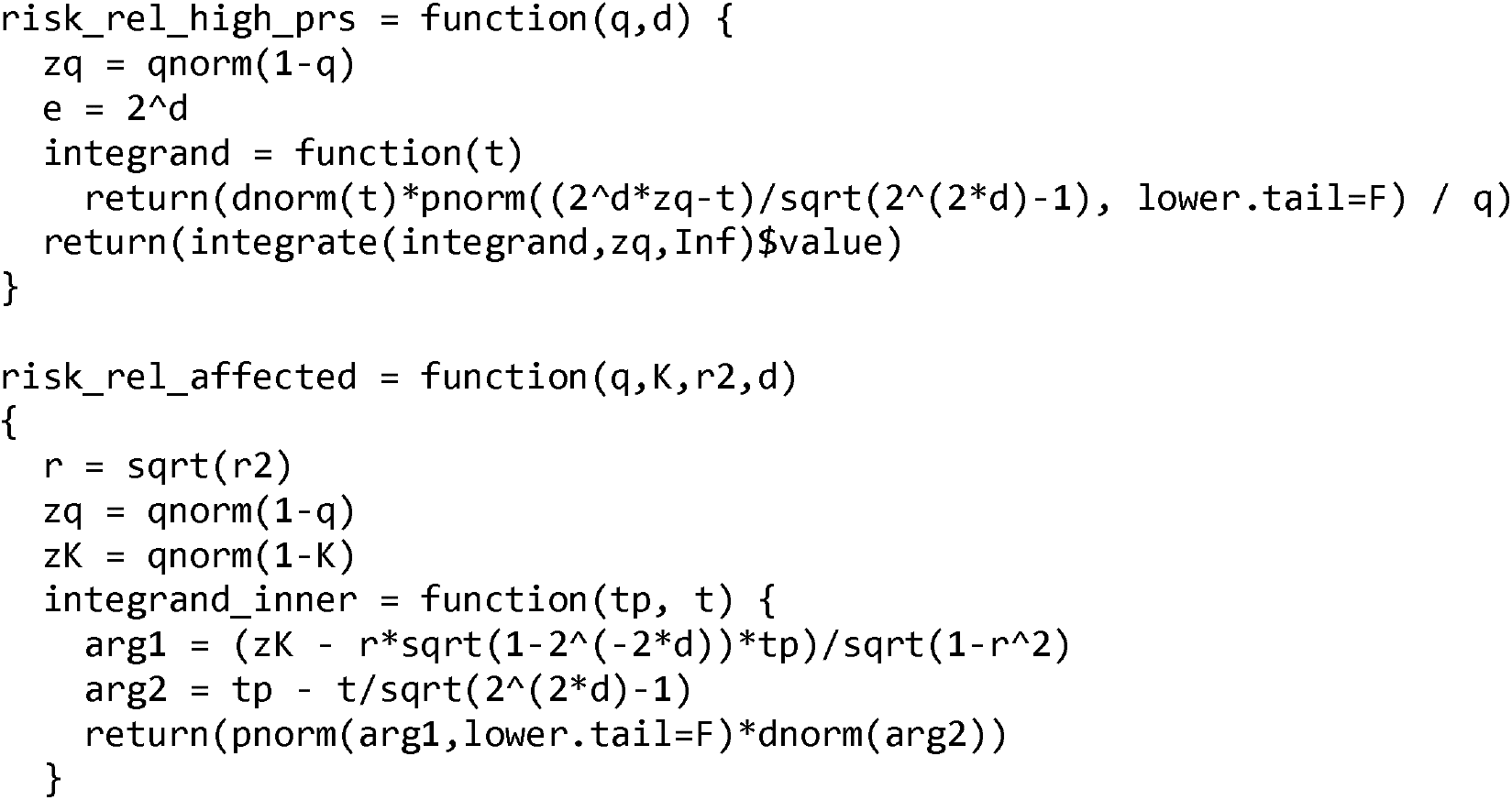
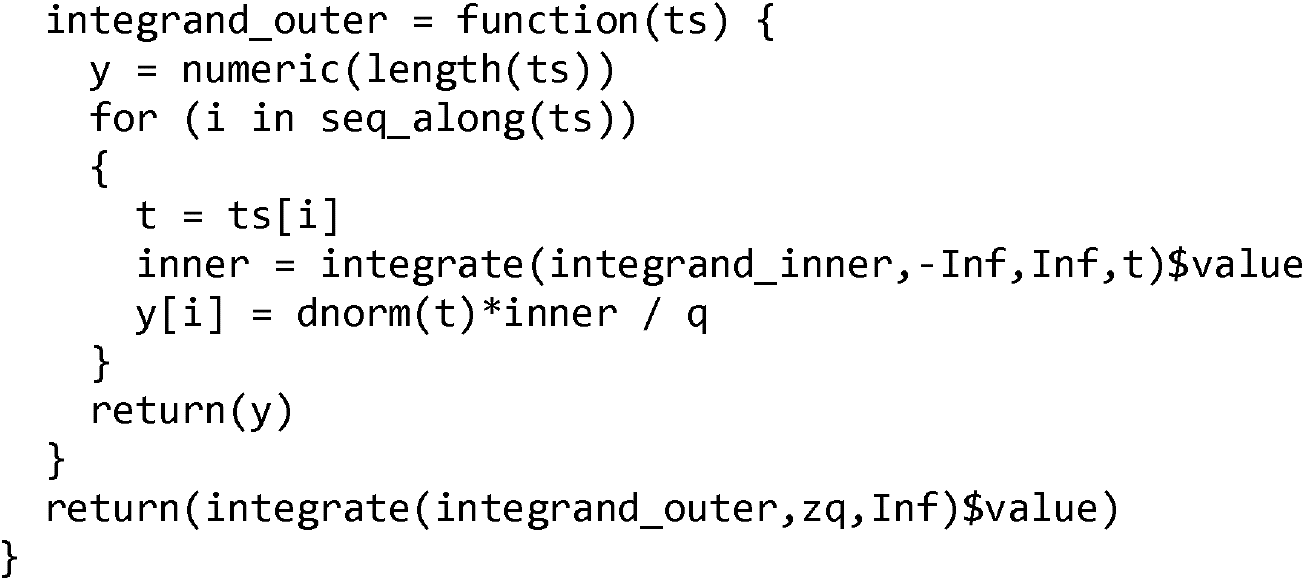

## Results and discussion

Reid et al.^10^ have first studied the correlation between the PRSs of relatives. They found that the correlation between the PRSs of siblings and second-degree relatives was ≈0.5 and ≈0.25, respectively. These correlations are naturally expected: based on standard quantitative genetic theory, the correlation between the genetic values of relatives is equal to their coefficient of relatedness^12,24^.

Next, Reid et al. have computed the empirical probability (across sib pairs in the UK biobank), that, given that the proband has PRS at the top *q* quantile, a sibling of the proband also has PRS at the top *q* quantile. Specifically, they considered four diseases (atrial fibrillation, coronary artery disease, diabetes, and severe obesity) and four values of *q* (1%, 5%, 10%, and 20%).

We used the liability threshold model to derive an analytical expression for the probability of the sibling of the proband to have PRS at the top *q* quantile (Methods; Eq. (5)). We compare our theoretical predictions to the empirical observations of Reid et al. in Figure 1A, showing excellent agreement. Our theory also predicts that the risk of the sibling is independent of the disease and the accuracy of the PRS, as empirically observed by Reid et al. Our figure also provides the expected risk for other values of *q* in the range 0-25%.

**Figure 1.**
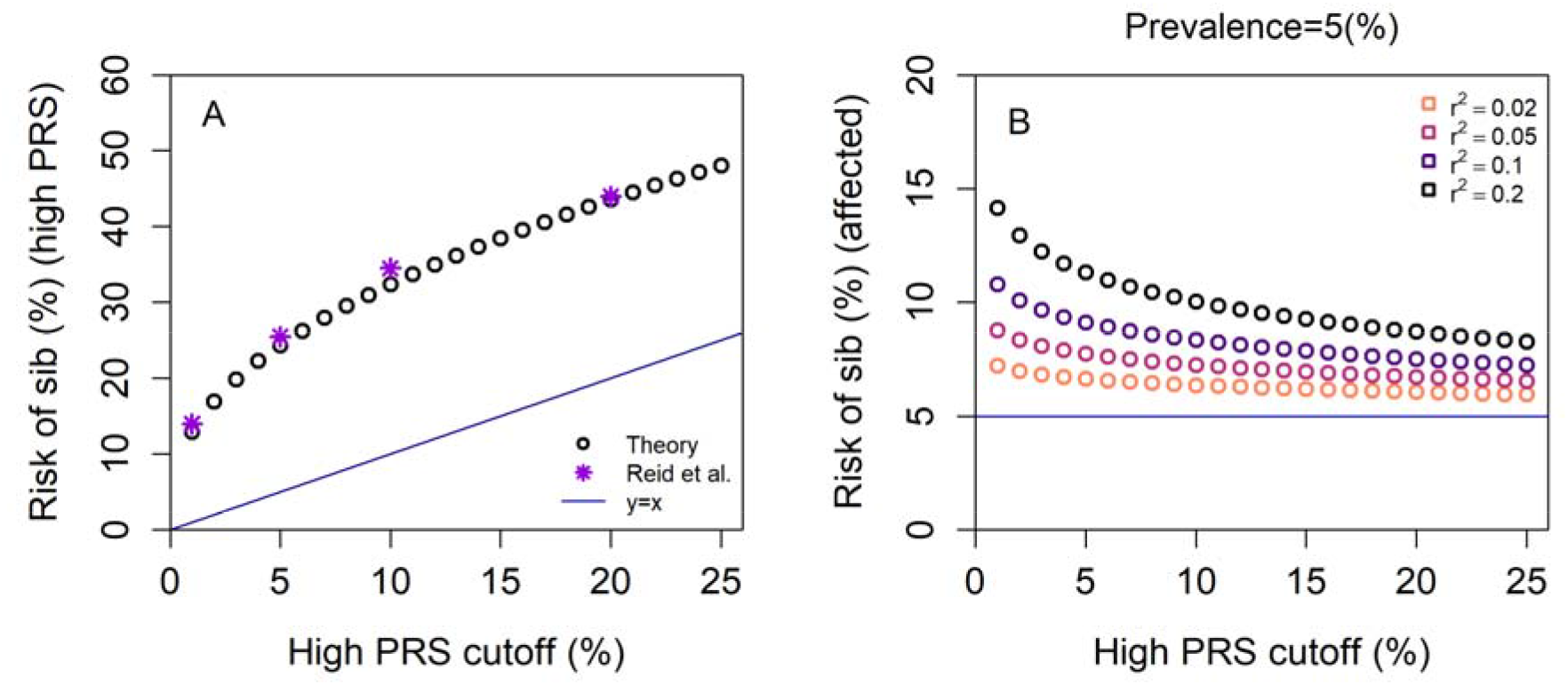
The expected risk of a sibling conditional on a proband having a PRS above a cutoff. We assume that the proband is known to have a PRS at the top q quantile of the PRS distribution. The cutoff percentile varies along the x-axis. In (A), we plot the risk of the sib of the proband to have a high PRS, defined using the same cutoff. The diagonal blue line is *y* = *x*, which is the risk for an unrelated individual. The circles are the theoretical probabilities we derived based on the liability threshold model, obtained by numerically evaluating Eq. (5). The violet stars correspond to the empirical values (mean across diseases) obtained by Reid et al. In (B), we plot the risk of the sib to be affected. We assume prevalence of *K* = 5%, and show results for multiple values of *r*^2^, a measure of PRS accuracy equal to the proportion of variance in liability explained by the PRS (legend). The horizontal blue line represents the risk of an unrelated individual (equal to *K*). The circles are the theoretical probabilities, obtained by numerically evaluating Eq. (11).

For studying the cost-effectiveness of cascade screening, it is necessary to estimate the risk of the sibling in the case screening is *not* applied. To this end, we need to compute the risk of the sibling to be affected, conditional on the proband having a high PRS. We used the liability threshold model to derive an analytical expression for this probability (Methods; Eq. (11)). We plot the results in Figure 1B, for a representative value of the prevalence (5%), and for four values of the proportion of variance explained by the PRS. As expected, the risk of the sibling is always higher than compared to a random individual from the population, and the risk increases with increasing accuracy of the score, and with a higher PRS of the proband (i.e., a smaller percentile used to define the top of the PRS distribution).

Full siblings are first degree relatives. To estimate the utility of cascade screening for more remote relatives of the proband, we extended our theory to compute the risk of a *d*-degree relative of the proband to have either high PRS (Eq. (16)) or to be affected (Eq. (23); Methods). While the resulting expressions are more complex, they are amenable to numerical evaluation. In Figure 2, we plot the risk of the relative to have high PRS (panel A) or to be affected (panel B) for *d* = 1,2,3,4,5 (*d* = 3 corresponds to first cousins and *d* = 5 to second cousins). As expected, as *d* increases, the risk of the relative to have high PRS or to be affected approaches that of the general population.

**Figure 2.**
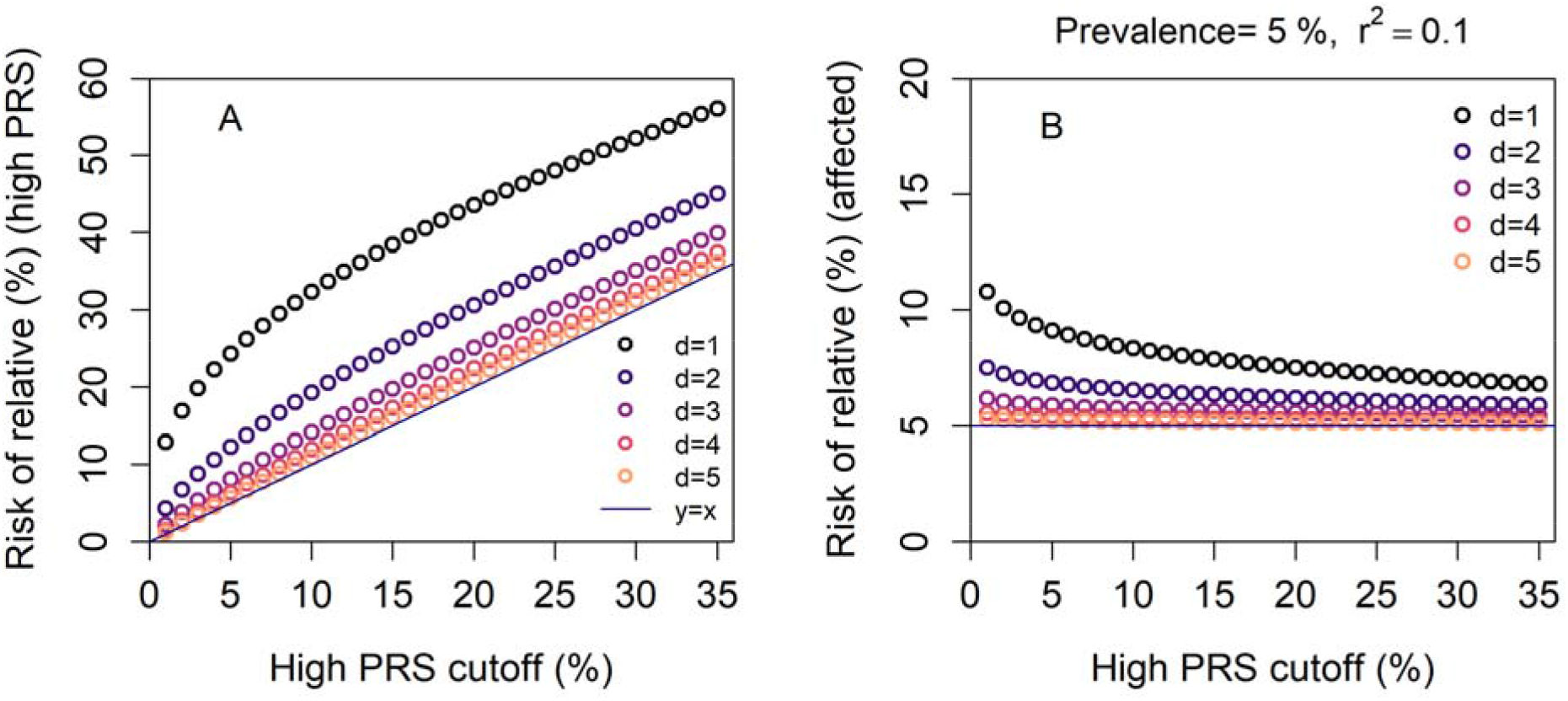
The expected risk of a d-degree relative of the proband. As in Figure 1, the x-axis represents the percentile cutoff used to define high PRS individuals. In (A), we plot the risk of the relative of the proband to have a high PRS, defined using the same cutoff. The diagonal blue line is *y* = *x*, which is the risk for an unrelated individual. The circles are the theoretical probabilities, obtained by numerically evaluating Eq. (16). Colors correspond to different degrees of relatedness *d* (legend). In (B), we plot the risk of the relative to be affected. We assume prevalence of *K* = 5%, and show results for multiple values of d (legend). The horizontal blue line represents the risk of an unrelated individual (equal to *K*). The circles are the theoretical probabilities (Eq. (23)).

Our simple R code will allow researchers to substitute any value for the high-PRS threshold, the accuracy of the score, the disease prevalence, and the degree of relatedness, in order to compute the expected outcomes of cascade screening in any setting of interest. Further conditioning on the disease status of the proband can be incorporated as in ^17,19^. We expect our results to form a necessary building block for future studies of the cost-effectiveness of cascade screening following a PRS test.

